# Habitat selection during dispersal reduces the energetic cost of transport when making large displacements

**DOI:** 10.1101/2023.11.25.568690

**Authors:** Tullio de Boer, Kennedy Sikenykeny, Brendah Nyaguthii, Damien R. Farine, James A. Klarevas-Irby

## Abstract

Dispersal is energetically costly. However, there is now growing evidence that dispersing animals can express distinct movement strategies that allow them to mitigate most of the energetic costs of displacing over large distances. Whilst to date we know that these strategies involve changes in how dispersing animals move, it is unclear whether these changes in movement are facilitated by other components of behaviour—namely changes in habitat selection. Here, we combine step-selection analyses with models of the energetic costs of movement to high-resolution GPS tracking data in terrestrially dispersing vulturine guineafowl (*Acryllium vulturinum*) to test the hypothesis that dispersing animals should select for habitats that facilitate more energetically efficient movements during dispersal. We find that actively dispersing individuals exhibit increased positive selection for open habitats, especially roads. We then show that moving along roads facilitates straighter, faster movements and results in a more than 33% reduction in the energetic cost of transport relative to other habitat types. Our results confirm that fine-scale differences in habitat selection expressed by dispersers facilitate more energetically efficient movement, expanding our understanding of how animals exhibit adaptive movement strategies across different axes of decision-making (e.g., where and how to move) to overcome ecological challenges.

## Introduction

Dispersal is a complex and often costly process. There are many factors that can drive individuals to move away from their natal habitats [1–3]. Individuals may need to move large distances to avoid inbreeding or competing with kin [4,5], to gain new mating opportunities [6], or to escape suboptimal environmental conditions [7,8]. The costs associated with making large displacements are illustrated by the many striking adaptations that animals and plants have evolved to enable long-distance movements, such as wings for flight [9–11], floating or rafting behaviours [12], and many mechanisms for seed movement [13,14]. In some species (e.g. many in the order Hymenoptera [15,16]), the traits associated with increased movement ability are expressed exclusively in the context of dispersal, highlighting that dispersal is a distinct life stage with its own trade-offs and adaptations. While many larger animals, such as vertebrates, cannot express extensive morphological plasticity across life stages (although there are clear adaptations, such as in wing morphology [11,17], to facilitate migration), there is nonetheless substantial scope for behavioural strategies to evolve to overcome the challenges that individuals face when dispersing.

For most species, dispersing requires individuals to make much larger displacements than they do throughout the rest of their lives [18]. It has long been assumed that these large displacements necessarily invoke an increase in energetic expenditure [18,19]. Logically, moving several times further than on an ordinary day [20,21] must consume more energy. However, studies spanning a range of terrestrial species—including bears (*Ursus arctos*) [22], lions (*Pantera leo*) [23], vulturine guineafowl (*Acryllium vulturinum*) [21], and elk (*Cervus canadensis*) [24] among others [25,26]—have revealed key changes in behaviour during dispersal that animals can employ to mitigate these costs. Actively-dispersing (transient) individuals were observed to achieve larger daily or seasonal displacements by increasing the speed and straightness of their movements. Doing so can increase energetic efficiency. This is because the non-linear relationship between movement speed and energetic expenditure [27] means that while moving faster uses more energy per unit of time, the corresponding time savings result in less energy used to cover a given distance relative to slower movement (i.e. greater energetic efficiency over space). Similarly, making straighter movements can also increase efficiency when making large displacements as it requires covering less overall distance than making more tortuous movements, and because making turns is also costly [28]. While few studies have quantified the relative energetic impact of these changes in behaviour (although see [26]), recent work in vulturine guineafowl has shown that these efficiency-driven behavioural strategies can allow individuals to achieve much greater displacements with only minimal increases in energy use relative to typical non-dispersal movements [21].

While increased movement speed and path directedness can result from behavioural changes, characteristics of the habitat, such as vegetation density, are also likely to determine the ability for individuals to express these movement characteristics [29,30]. For example, dispersing African wild dogs (*Lycaon pictus*) typically make larger or more directed movements in open (e.g., grassland) versus closed (e.g., woodland) habitats [31], a pattern that is mirrored in several other species (e.g., elk *Cervus canadensis* [24], and red squirrels *Sciurus vulgaris* [32]). During foraging movements, olive baboons (*Papio anubis*) preferentially move along roads in the morning and evenings, when commuting between their sleeping site and distant foraging areas [33]. However, like many aspects of the transient phase of dispersal [34], changes in habitat selection during dispersal (i.e. relative to habitat selection within a settled home range) have been generally overlooked as a means for animals to mitigate the movement costs associated with dispersing. In particular, when faced with higher energetic costs to travel large distances, animals should be expected to selectively use habitats that facilitate faster, straighter, and thus more efficient movements.

Animals frequently express flexibility in habitat use according to their current needs. For example, many animals move from open foraging areas to shade or cover to maintain thermal homeostasis [35] or to avoid exposure to predators [36] at times of day when hyperthermia or predation risks are highest. In the context of dispersal, and especially in terms of making large displacements, we expect that transient animals should also adjust their habitat preferences by disproportionately selecting those that maximise energetic efficiency associated with movement. Due to reduced barriers to movements and increased navigational ease arising from greater visibility, we expect that dispersers should preferentially move through more open habitats. However, in social species, open habitats may also provide benefits in terms of increasing the likelihood of detecting conspecifics [37]—whether to facilitate grouping [38] or for avoiding aggression from territory owners [39]. Thus, while selectivity for open habitats during dispersal has been observed in several species, it has yet to be confirmed whether such preferences translate to increased energetic efficiency.

Here we use high-resolution GPS tracking in a wild population of vulturine guineafowl to assess (i) the habitat selection behaviours of dispersing individuals, and (ii) quantify whether the habitat features that dispersing vulturine guineafowl select for correspond to a reduction in the cost of transport—a measure of efficiency capturing the energy consumed to move a given distance— associated with making large displacements. Vulturine guineafowl are almost exclusively terrestrial and live in large groups varying in size from 13 to 65 individuals [40]. When dispersing, individuals can walk up to 15 km per day, doing so mostly alone [21] while traversing a mixture of savannah scrub, open grassy areas, and roads—all habitat features that are similar to those present in the home range of their natal group. Previous work has typically compared the dispersal period to the behaviours of the same individuals prior to dispersing (e.g. [41,42]). However, in many systems, such as buzzards (*Buteo buteo*, [43]) and wolves (*Canis lupus*, [44]), the timing of dispersal is also linked to changes in ecological conditions [45,46]. In the vulturine guineafowl, dispersal is linked to periods of greater rainfall [47], which is also likely to alter habitat conditions. This requires making several contrasts in order to confirm that habitat selection during dispersal reflects the energetic cost—and savings therein—of key habitats.

Because vulturine guineafowl remain in temporally stable groups when not dispersing [48], we can directly contrast the habitat selection behaviours that individuals express during dispersal to those that they would have expressed (as part of their group) if they were not dispersing, making this system ideal for analysing habitat selection (and its consequences) during dispersal. Further, because dispersers express a bimodal distribution in terms of displacement during the transient phase of dispersal [47], we can further examine whether changes in habitat selection are specifically linked to periods when they are actively dispersing (i.e. making larger displacements) versus interspersed periods when they are prospecting locally for groups to join. Guineafowl are also a model system for physiology, with detailed data available on their energetic expenditure when walking at different speeds [49]. Previous work combining physiological models with second-by-second movement data has revealed that, when dispersing, individuals experienced only an average 4.1% increase in the daily cost of transport relative to their normal daily ranging behaviours, despite achieving an 33.8% increase in displacement [21]. In this study, we confirm the prediction that the increased energetic efficiency expressed during dispersal is facilitated by moving through open habitats. We do so by (1) showing that vulturine guineafowl are more likely to select for open habitats (e.g. grassy areas and roads) during dispersal relative to non-dispersal movements, (2) by revealing that walking through these habitats corresponds to a lower average cost of transport, and (3) showing that this benefit comes from dispersers being able to express faster, straighter movements (as opposed to benefitting from these habitats having less inclined or rugged topography).

## Materials and Methods

### Overview

Our study consisted of four components (Fig. 1).

1. Field Data Collection: We fitted GPS tags to subadult vulturine guineafowl and to adults from their natal group in order to collect high-resolution GPS data on the position of each bird.
2. Environmental Data Collection: We used remote sensing to extract features of habitats through which both resident and dispersing individuals moved.
3. Step-Selection Analysis (Fig. 1b: We used a step-selection analysis (SSA) [50] framework to infer the habitat selection behaviours of individuals, which we split across three categories: actively-dispersing transience movements, local movements while transient, and the non-dispersal movements of resident adults (date-matched to periods of dispersal).
4. Estimating Energetic Efficiency in Different Habitats (Fig. 1c): We combined published relationships of energy expenditure as a function of movement speed and incline with high-resolution (1 Hz) GPS data from individuals to quantify the average energetic efficiency of movement (the cost of transport for a given net displacement) that they expressed when moving through each habitat type.

**Figure 1.**
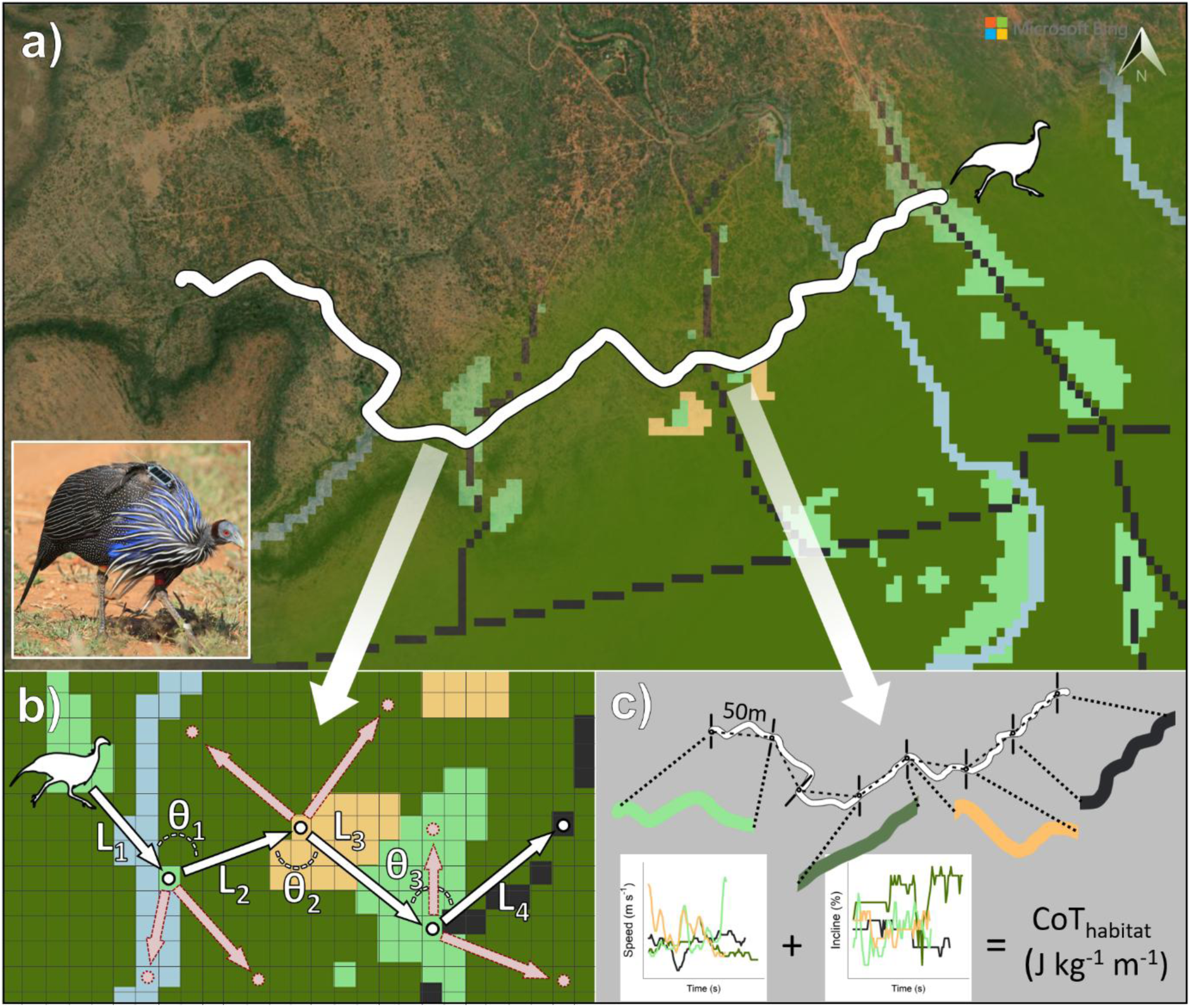
Overview of study methods and analyses. (a) GPS tracks (white) of vulturine guineafowl movement were collected from wild birds (GPS tagged guineafowl, inset, photo by [author anonymised]) in Laikipia, Kenya and matched to rasters of underlying habitat features, such as roads (black dashed lines), acacia scrub (dark green) and more open habitats like glades (light green) and bare soil (orange). (b) To quantify habitat selection according to different stages (active dispersal, local movements, resident birds), we conducted a step-selection analysis, using observed steps from the GPS track taken over 5-minute intervals (white arrows and points) and alternative steps (dashed red arrows and points, generated based on observed step length (L) and turning angle (θ) distributions). (c) To quantify the energetic consequences of moving through the selected habitats, we extracted track segments consisting of 50-meter net displacements which occurred entirely within a given habitat type and estimated the energetic cost of transport based on the speed and incline of movement from GPS data collected at 1Hz [21]. Satellite imagery generated using Bing Maps, Maxar Technologies (accessed May 2025).

We performed all analyses in R version 4.3.1 [51].

### Study System

This study was conducted at the Mpala Research Centre in Laikipia county, central Kenya (0.29120 N, 36.898670 E). The research centre is part of the Mpala Conservancy, which covers an area of roughly 200 km^2^ and is part of the Greater Ewaso Ecosystem [52]. However, because vulturine guineafowl disperse outside of Mpala, our study extends beyond these boundaries to cover approximately 769 km^2^. Within this broader study area, the landscape contains a heterogeneous mix of habitat, which can broadly be divided into three major types. In the more elevated areas, we find a type of clay vertisol commonly known as “black cotton” soil [53] and its dominant tree species the whistling thorn (*Acacia drepanolobium*), and which is a habitat largely avoided by vulturine guineafowl. On the lower part of the escarpment typical “red soils” which contain a mixture of Acacia scrub and open glades as well as riverine habitats with fever trees (*Vachellia xanthophloea*) are present [54], and vulturine guineafowl predominantly reside on these red soils. Both major habitat types are crossed by numerous roads and smaller vehicular tracks used for research and management within Mpala and surrounding reserves.

Our study on vulturine guineafowl has been running since 2016 [55]. The present study, however, relies on a shorter, more focused, period—September 7th to November 25th, 2019—during which we had fitted GPS tags (see below) to female vulturine guineafowl that were still in their natal group. Previous work using this GPS-tagged cohort, together with field observations, has suggested that dispersal is exclusive to females, typically solitary, and occurs around two years of age, when individuals are physically mature [21]. However, because dispersing guineafowl do not form new groups but disperse between existing groups [55], they often fail to settle into a new group (due to social resistance, *sensu* [56]) and return to their natal group. They can remain in their natal group for several months before trying again. There are typically two dispersal seasons per year, corresponding to the major and minor wet seasons, with our study corresponding to the minor wet season of 2019. As the study cohort is part of a larger, GPS-tracked population, we studied the movements of both dispersing individuals and non-dispersing resident members of their natal group during the same time period. In doing so, we can compare movements under the same environmental conditions and assess whether there are differences in the types of habitats selected. Notably, group movements in this species emerge from democratic processes (i.e. following the preferences of the majority), where any given individual can initiate movement in a preferred direction [57], facilitating comparisons of habitat selection behaviours between dispersing individuals and residents.

### Field Data Collection

Birds were captured using baited walk-in traps, allowing us to trap the entire group at once. All individuals were marked on their legs with a uniquely numbered stainless-steel ring and a distinct combination of four plastic colour bands, allowing individual identification in the field. Within each group, we selected individuals (typically 2-4 resident adults of both sexes) to be fit with a GPS tag (15g Bird Solar, e-obs GmbH) using a backpack-type harness (see [40] for more details). For the purposes of this study, 18 dispersal-age females from four social groups were fitted with GPS tags, which were complemented by 13 GPS-tagged resident adults (9 males, 4 females). The combined weight of backpacks, tags, and leg markings is less than 3% of birds’ body weight.

Tags were programmed to record fixes from 06:00 to 19:00 local time (corresponding to daylight hours when the birds are active). Fix data (date, time, and location) were recorded at two resolutions: continuous high-resolution (i.e., 1Hz) data was recorded when tag batteries had a high level of charge (up to 4.5h continuous recording, approximately every second to third day), and at a lower resolution—collecting a 10-second burst of 10 fixes every five minutes—when battery levels fell below the high-resolution threshold. He et al. [58] provide further details on the strategy behind this sampling design. Data were remotely downloaded from tags over a secure VHF connection at least every other night after birds had entered their roosts. For transient individuals, we downloaded data every morning and evening once we first detected birds departing from their natal group. In case of morning download, we traversed as much of the landscape as possible to opportunistically detect and download data from actively dispersing birds (additionally aided in obtaining a bearing for relocating them in the evening). Some birds ultimately dispersed beyond the area which we could access their GPS tags for download (although three such individuals were eventually re-encountered after several months and their accumulated date was recovered). GPS data were uploaded to the Movebank repository (https://www.movebank.org) for remote storage and prepared for analysis in R through the move package [59].

### Dispersal

We defined three stages of dispersal (departure, transience, and settlement) based on key features of our tracking data. Birds were considered to have departed (i.e., to have started dispersing) on the first date when their GPS tracks diverged from those of the resident adults in their natal group. Dispersers were subsequently considered to have settled in a new area after using the same roosting site for 14 days (settlement was back-dated to the first date after they entered the new roost). Where possible, departure and settlement events were confirmed by visual observation of the GPS tracks (i.e. we observed no further large displacements by a given individual later than the 14-day cutoff) and field observations of guineafowl groups (confirming that dispersers were part of a new group and not ranging alone). All GPS data between the departure and settlement dates for each individual were then treated as transience. In 11 cases, we were unable to record settlement—either due to dispersers failing to settle and returning to the natal group after a period of several weeks (3 events), due to predation during transience (1 event), because the dispersing individual moved far enough from the study area that the tag could not be relocated for downloading data (6 events), or due to the harness failing and the tag falling off (1 event). In these cases, all transience data up to the available end point were used. As this study focused on the habitat selection behaviours of transient birds, all pre-departure and post-settlement GPS data were excluded from our analyses. In order to facilitate comparisons to non-dispersing residents, we included date-matched GPS data for each disperser from all available resident adults of each dispersers’ natal group.

Dispersal movements while transient are not uniform over time in vulturine guineafowl [47,60], but differentiated into two distinct daily patterns. The first involves birds making large roost-to-roost displacements, corresponding to days of active dispersal characterized by extended periods of long, straight movements across the landscape. The second involves movements that take place over much smaller local scales (often without changing roosts from morning to evening) with more tortuous paths. Field observations suggest that local movements are the result of dispersers temporarily joining and moving with groups other than their natal group for one or more days before resuming large displacements (first category). As such, we split each day of data from each disperser into two categories based on the length and straightness of each day’s movement path. Specifically, an individual was considered to have engaged in large, active dispersal movements if the distance between consecutive days’ roosts exceeded 1500 m, or else if the roost-to-roost distance was greater than 1200 m and the ratio of this distance to their total daily track length (i.e., a straightness index [61]) was greater than 0.3 [47]. Sub-categorizing the movements of transient individuals in this way allows us to better disentangle the habitat selection behaviours of actively dispersing birds from the overall effect of being transient, and compare each to the behaviours of non-dispersing residents. See Supplementary Materials for description of total GPS dataset, including distributions across stages of dispersal and step parameters.

### Environmental Data Collection

#### Generating Habitat Raster Layers

To measure and define the overall habitat composition of our study area, we downloaded satellite imagery from Planet Explorer by Planet Labs [62]. The boundary of our satellite data was selected such that all GPS tracking data fell inside a bounding box, which was then used to build the base geometry for the search and download of suitable satellite imagery. We selected the PlanetScope Scene imagery (hereafter, “satellite imagery”) from the 3rd of November 2019 (resolution: 3 m, area coverage: 94%), as it best satisfied our criteria (maximum area coverage, no cloud coverage) within the period when we recorded the majority of our recorded dispersal events. Satellite imagery was then prepared and processed for analysis in ArcGIS’ ArcMap 10.6.1. [63]. The satellite images were georeferenced, classified, and projected to WGS84 UTM Zone 37N (EPSG: 32637). We manually extracted three raster layers (Fig. 2) capturing roads, water bodies, and habitat type (the latter being classified into 5 types).

**Figure 2.**
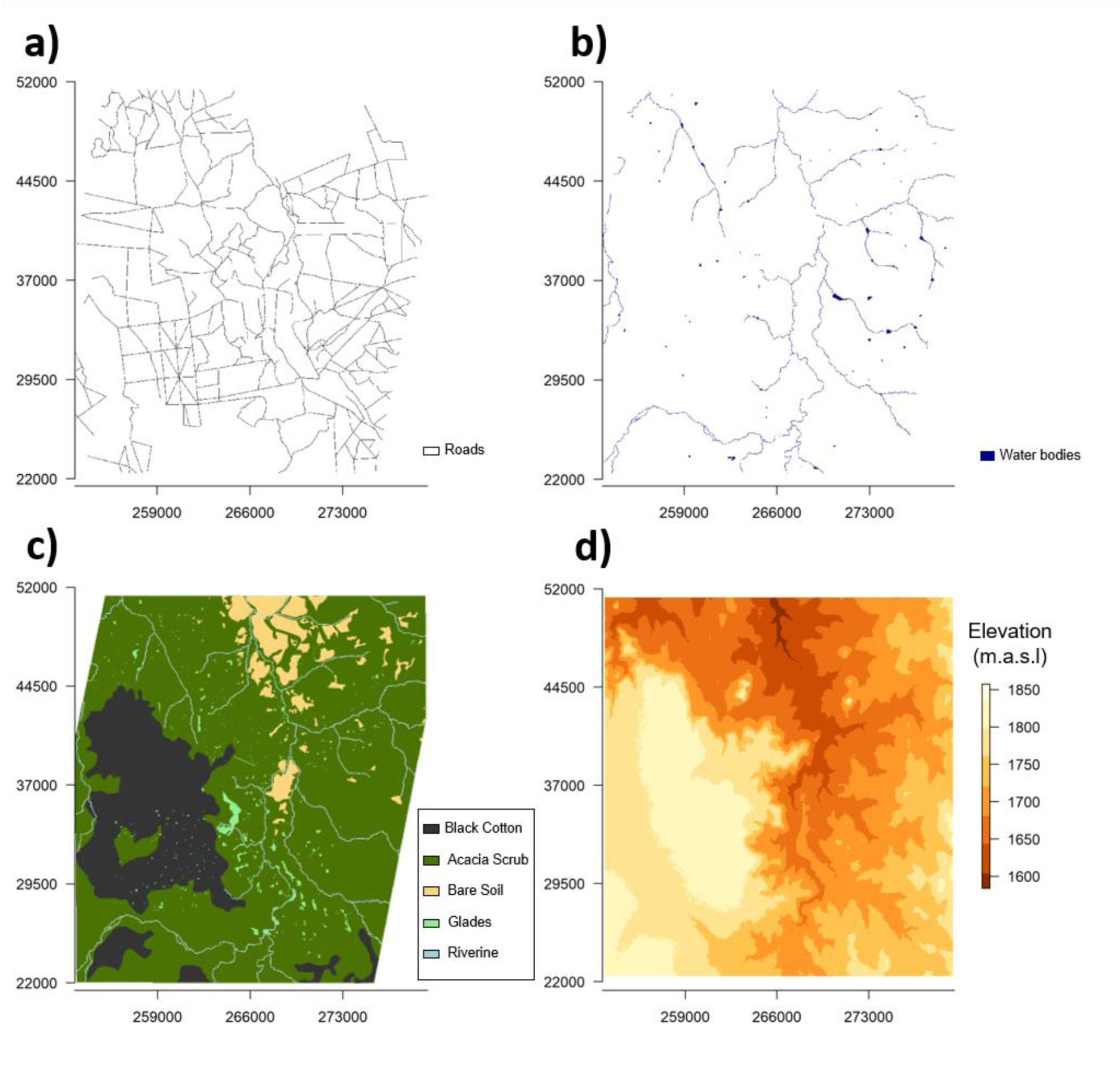
Raster layers of habitat covariates across our study area. Habitats were classified according to the presence of roads (a) and waterbodies (b), as well as by predominant cover type (c) and elevation (d) in meters above sea level. Axes show Universal Transverse Mercator (UTM) coordinates.

Roads and water bodies were both defined as binary variables, meaning that all classified pixels contained either presence (1) or absence (0). As no road network layer was available for our study area, we manually marked roads as polylines. With our spatial resolution of 3 m, it was not always easy to distinguish between smaller or older/abandoned roads and animal paths. Thus, we opted for including roads that were distinguishable through visual inspection of our satellite data and Google Earth. As most roads are between 4 and 10 m wide, we defined roads as polygons which extended 15 m to either side of the centre of each polyline. All roads were rural gravel roads, with no heavy-use roads present in the study area.

For water bodies, we followed the same approach as for roads to obtain polylines for rivers (buffering 15 m on each side). Ponds, dams and small lakes were drawn as polygons and merged with the river features. Water pixels could be distinguished from other landscape features based on their reflection in false colour infrared or in the NDVI. The main component for the water bodies layer in our study area is the Ewaso Ng’iro river and some of its side channels. There are also a few ponds, small lakes, and dams; the largest being a dam on the nearby Ol Jogi reserve with an area of 0.3 km^2^.

Utilizing the post-processed satellite imagery, we categorized habitats into five cover types. Categorization was done using the false colour infrared band combination and was then compared to Google Earth scenes with higher spatial resolution and true colour. Black cotton soils, due to their dark colour, were easily detectable as dark tones in the false colour infrared spectral combination. Glades (open grassy areas) appeared in vivid red in the same band combination, due to the higher chlorophyll content found in the plant species on open glades (given that green reflectance shows as red in false colour infrared) compared to surrounding vegetation. Bare soil represents sandy patches with little to no vegetation and was distinguishable through its white pixel colouring. The predominant habitat cover in our study area was dense acacia scrub, which presented as a dark reddish colour, due to its lower chlorophyll content compared to glades. Finally, we classified a 30 m buffer zone along marked rivers as Riverine habitat.

#### Generating the Topography Raster Layer

To quantify topography, we used a digital elevation model (DEM) from the Open Data Site of the Regional Centre for Mapping of Resource for Development (opendata.rcmrd.org) with a spatial resolution of 30 m. We then calculated a Terrain Ruggedness Index (TRI) which quantifies average topographic heterogeneity [64]. TRI values were calculated by taking the DEM values from a centre cell and the differences to its surrounding eight neighbour cells (we ground-truthed the DEM by measuring elevation at 24 locations in the study area). The differing values to each of the eight neighbour cells were then squared, summed up and subsequently square rooted [64], which effectively quantifies the topographical heterogeneity in a given area with higher values corresponding to more rugged or uneven terrain.

### Step Selection Analysis

We used the ‘amt’ package [65] in R to conduct a step selection analysis (SSA) to assess how landscape features influence where vulturine guineafowl move according to different stages. This type of analysis uses positional data, and movements from one location to another, to generate movement steps. Each of these observed steps (i.e. two consecutive relocations in our GPS data) was then matched with a chosen number of alternative steps [50] that represent plausible alternative locations in which the individual could have moved (given the properties of its movements). The environmental features were then compared between the observed and alternative steps to identify which were over-represented in the observed steps.

#### Temporal Discretization of Observed Steps

In order to maximise the continuity and coverage of our data, we opted to divide our data into 5-minute steps (combining continuous, 5-minute data with sub-sampled 1Hz data). After calculating step lengths (net displacements) for each 5-minute interval, we removed all steps shorter than 10 m to facilitate our focus on where an animal moved rather than on the timing of movement [33]. In this way, the resulting model is less affected by periods of little or no movement. We then marked the next time point as the start of a new step, and repeated the process for the rest of the track. From an initial dataset of 132,959 5-minute GPS intervals, our final dataset consisted of 61,435 observed steps greater than our 10 m threshold.

#### Generating Alternate Steps

For each observed step, we recorded the true step length (in meters) as well as the turning angle between consecutive steps (in radians). The distributions of these values (Supplementary Fig. S1) were subsequently used to generate a series of randomized alternative steps for our SSA. For each individual and for each category of movement (i.e. active dispersal or local movements) they expressed, we fit a gamma distribution to the observed step lengths and a von Mises distribution to observed turning angles [66]. For each observed step, we generated 20 alternative steps based on the resulting individual-and-category-level distributions of step lengths and turning angles.

#### Linking Steps to Habitat Features

Using the ‘raster’ package [67] in R, we combined our dataset of observed and alternate steps with our environmental covariates. We first imported our prepared environmental raster layers (i.e. presence/absence of waterbodies and roads, habitat cover type, and a terrain ruggedness index) and then projected our step locations onto them. For each step, we extracted the relevant environmental covariates from the raster cell which contained the endpoint of that step.

#### Fitting a Step Selection Function

Using ‘amt’, we fit a wrapper function of a conditional (fixed-effects) logistic regression for matched case-control (observed vs. alternative step) data with the R package ‘survival’ [68]. We fit the conditional logistic regression using the same routine as a Cox Proportional Hazards model [65]. Each observed step and its matching 20 alternatives formed a stratum together. We fit two models, which produce comparable selection coefficient estimates, but which evaluate significance based on differing criteria (see Supplementary Tables S1-S2 for detailed model formulas). The first model was designed to assess whether the selection behaviours exhibited within each movement stage (i.e. resident, local-transient or actively-dispersing transient) were significantly positive or negative (representing selection or avoidance, respectively). This first model was fit to an interaction term capturing each of our environmental covariates—water bodies, roads, cover type (with acacia scrub as the reference level), and terrain ruggedness— interacting with a three-level categorical factor representing movement stages (where residents were set as the reference level). The second model assessed whether dispersing birds exhibited selection behaviours which differed significantly from those of non-dispersing residents (i.e. regardless of if the overall selection strength was positive or negative). This second model featured a standalone term for each of our environmental covariates as main effects, in addition to the interaction term for each environmental covariate with movement. We also constructed a second pair of supplementary models (see Supplementary Tables S3-S4) with only two levels of interaction (transient and resident, irrespective of the type of daily movement during dispersal) to enable comparisons between the present study and other systems where analyses are more typically divided based on whether or not individuals are dispersing. Models estimate habitat selection as the relative strength of selection (RSS) for a given habitat feature, corresponding to the probability of selecting for a location which corresponds to a given feature relative to the reference level (assuming both options are equally available).

Non-dispersing residents made no observed steps into areas with black cotton or bare soils (Supplementary Table S5), and initial models showed no strong evidence for selection or avoidance of these habitats by dispersers. As such, we made the post-hoc decision to remove all steps (observed and alternate) in black cotton and bare soils, and re-fit all of our step selection models without them. This avoided spurious estimates of selection where residents were used as the reference category for movement stage but where they were not represented in the data for these habitat types.

### Estimating Energetic Efficiency in Different Habitats

We conducted a separate analysis to generate estimates of the efficiency of movement (here the energetic cost of transport) for each habitat and stage of movement (i.e. non-dispersing residents, localised transient movements, or transient birds making large displacements). Because many dispersing individuals moved through areas that were not used by any other GPS-tagged bird, we could only calculate efficiency metrics for observed movements (i.e. we could not infer layers of energetic landscapes). Thus, we ran this analysis independently of the step-selection analysis, generating a per-habitat measure of cost of transport.

#### Spatial Discretization of Net Displacements

The cost of transport is a measure of efficiency that describes the energy used to displace over a given distance. We obtained the cost of transport by first dividing our high-resolution (1Hz) movement tracks into net displacements of a fixed size (here 50 m). These net displacements were extracted using first location from each high-resolution GPS track and, for each successive time point (current location of the bird) calculating the net displacement (or Euclidian) distance to the first location, until a threshold radius of 50 meters was reached, which we marked as the end of the displacement. We then marked the next time point as the start of a new displacement, and repeated the process for the rest of the track. The resulting dataset contained relocations approximately every 50 m along with data on the total movements between each point. Although our threshold was set at 50 m, the true distance of a displacement often slightly exceeded this (owing to the temporally discrete sampling of our tags), and so the true distance of each net displacement was used for the calculations of the energetic cost of transport (i.e., per-unit-distance energy expenditure, in Joules kg^-1^ m^-1^) that followed.

#### Linking Energetic Costs to Habitat Features

We first sub-sampled our dataset of 50 m net displacements for all instances in which the start and endpoint of a given net displacement were within the same habitat or landscape feature, operating under the assumption that individuals moved through continuous habitat between the two points. If the movement took place on a road, we defined the category as “road” for this analysis, independent of the habitat cover that the road crossed. For all other categories, the cover type of the movement was used as the categorical label.

For each high-resolution fix within a given net displacement, we derived the metabolic cost of movement (in mL O2 kg^-1^ s^-1^) at each second based on laboratory studies describing how the metabolic costs of movement vary across different controlled movement speeds (m s^-1^) and inclines in the morphologically-similar, closely-related helmeted guineafowl (*Numida meleagris*) [49]. When stationary, guineafowl exhibit a flat rate of oxygen consumption of 19.1 mL O2 kg^-1^ min^-1^. When moving, oxygen consumption increases linearly with speed (v), where the slope and intercept of that relationship vary depending on the incline at which birds are moving [49]. These formulae take the form of VO_2_ = 24.0v + 27.2 on level terrain, VO_2_ = 30.7v + 27.6 at 10% incline, and VO_2_ = 47.7v + 21.3 at 20% incline [49], where V0_2_ is the per-minute volume of oxygen consumed in mL O_2_ per kilogram of body mass (mL O_2_ kg^-1^ min^-1^). Some energetic savings can be realized during downhill movements, but these are often offset by increased muscle load to absorb additional impacts [69], leading to either minimal savings (e.g. [70]) or even slight increase the cost of transport (e.g. in barnacle geese *Branta leucopsis* [69]). As there are no studies showing the specific impact of downhill movement on guineafowl, we assumed any downhill movements followed the same cost relationship as those on level terrain.

In order to apply the correct cost formula to each second of data within a given displacement, we first determined whether an individual was actively moving. Movement was assigned according to the outputs of a 4-state Hidden Markov Model (constructed using the R package depmixS4 [71]) parameterized on 10-second step lengths and turning angles. We used a 4-state model because this was previously found to better capture movement states (and, in particular, to differentiate a non-moving state) than models containing fewer states [21]. When moving (i.e. assigned to states 2-4), we then calculated the degree of the slope experienced based on the relationship between the individual’s movement bearing and the aspect of the underlying incline (terrain slope and aspect were derived using the DEM elevation model above), following the formula θ^′^ = tan^−1^(tan(θ) · cos(Δψ)) where θ^′^is the slope experienced by the individual, θ is the slope of the inclined plane, and Δψ is the angular difference between the aspect of the incline and the individual’s bearing. Experienced slope values were transformed into percent grade by the formula *PG* = tan(θ′) ∗ 100 where *PG* is the percent grade of an incline corresponding to the slope experienced by an individual (in degrees). Experienced incline values were aligned with formulas for oxygen consumption by rounding the percent grade at each GPS fix the nearest 10^th^ percentile. Individuals were considered as moving on level terrain when resultant value was less than 5% (i.e. rounded to 0), were considered as moving at 10% incline when the rounded value was between 5% and 15%, and followed the formula for a 20% incline when resulting values were ≥15% (no points corresponded to an experienced grade >= 30%). Per-second measures of oxygen consumption were then calculated based on whether and individual was stationary or moving and, if moving, according to the speed and underlying incline. These measures were then transformed into units of energetic consumption (Joules kg^-1^ s ^-1^) using a conversion factor of 20.1 J mL^-1^ O2 [49,72].

#### Estimating the Energetic Cost of Transport Within Each Habitat Type

To calculate the cost of transport for each 50m net displacement, we summed the total per-second energetic costs within that displacement and then divided this sum by the true net displacement distance. The effect of habitat type on the energetic cost of transport was calculated by fitting three linear models where we modelled the cost of transport for each habitat type using a subset of the data from each movement stage (i.e. one model each for residents, local transience, and large transience displacements; full details in Supplementary Table S6). An additional post-hoc test showed no significant pairwise contrasts among non-road habitat variables in any of the three models, and so we only report estimated cost of transport values relative to roads (as modelled) here.

## Results

### Dispersers Differ in Habitat Selection to Residents

We found distinct differences in habitat selection across different stages of movement (i.e. within-stage selection behaviours for residents and transient birds moving over local or large scales), and by contrasting the two patterns expressed by transient birds against residents’ behaviours. Non-dispersing residents expressed significant positive selection for glades (Relative Selection Strength [RSS] = 1.382), significant avoidance of roads (RSS = 0.942), riverine cover (RSS = 0.942), and rugged terrain (RSS = 0.908), and no selection for or avoidance of waterbodies (Fig. 3a; see Supplementary Table S1 for full results). Actively dispersing transient birds (i.e. when making large displacements) expressed significant positive selection for roads (RSS = 1.166) and glades (RSS = 1.636), significant avoidance of riverine cover (RSS = 0.693) and rugged terrain (RSS = 0.926), and no selection for or avoidance of waterbodies (Fig. 3a; Supplementary Table S1). During smaller, local movements, transient birds exhibited significant positive selection for glades (RSS = 1.719), significant avoidance of riverine cover (RSS = 0.724) and rugged terrain (RSS = 0.916), and neither selection nor avoidance towards roads or waterbodies (Fig. 3a; Supplementary Table S1).

**Figure 3.**
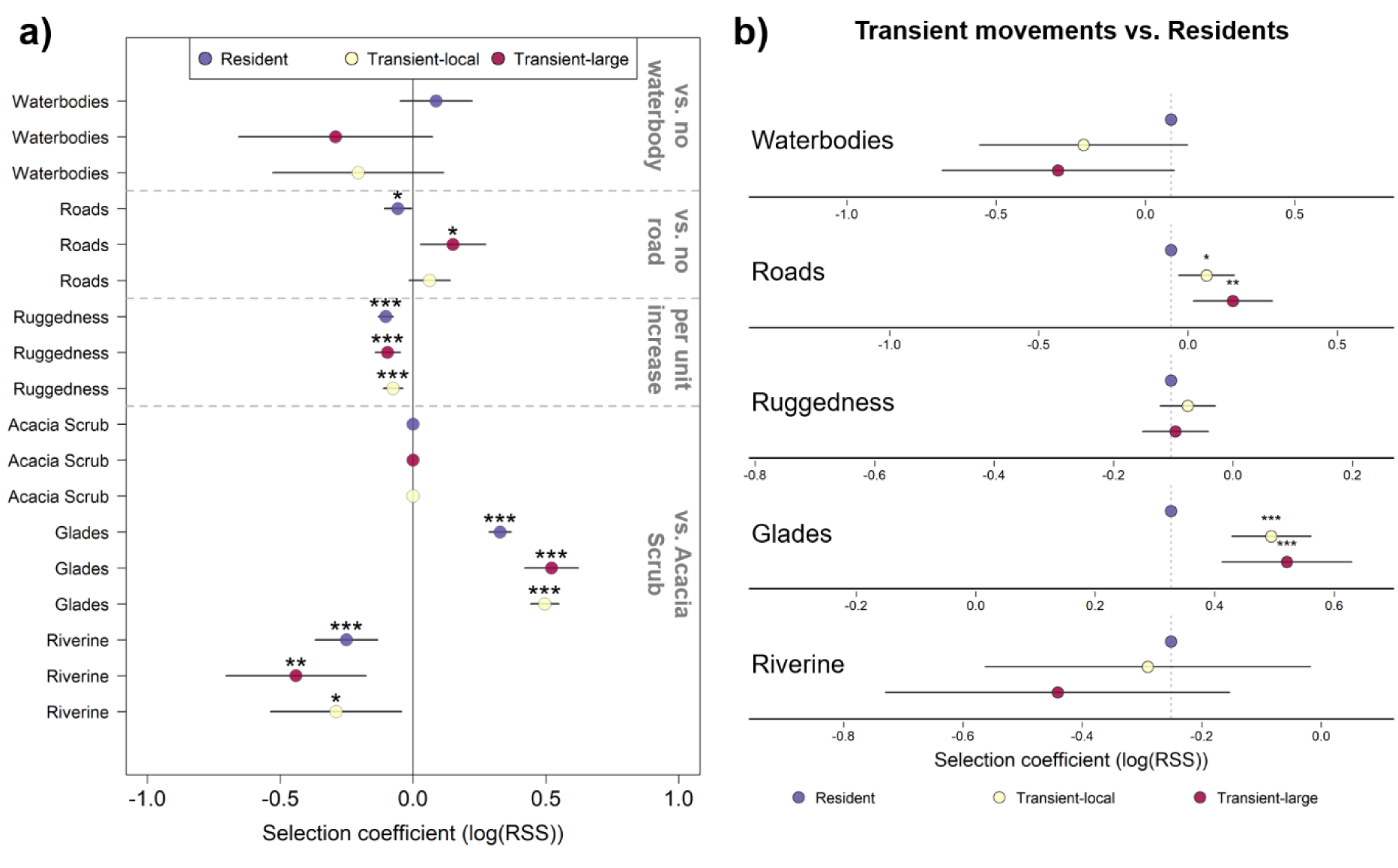
Relative Selection Strength (RSS) for different habitat features differs between residents and transient dispersers. (a) Strength of selection within a given stage of movement (non-dispersing residents—purple; transient birds making small, local movements—white; actively-dispersing transient birds making large displacements—red). Points represent log-transformed estimated relative selection strength (RSS) values for a given habitat feature relative to a reference category (grey text, right; separate raster layers denoted by dashed grey lines). Black bars represent 95% confidence intervals, indicating strength of habitat selection for (where log(RSS) > 0) or avoidance of (where log(RSS) < 0) a given habitat type. Asterisks above a point indicate the statistical significance (*** *p<0.001*, ** *p<0.01*, \**p<0.05)* of a given estimate. Full model results and formula in Supplementary Table S1. (b) Selection strength by transient birds relative to residents. Selection coefficients and confidence intervals same as in panel a, with statistical significance evaluated base on relative difference to the reference category (residents— marked with purple points and vertical dashed lines) for each habitat feature. Asterisks above points (\*\*\**p<0.001*, ** *p<0.01*, \**p<0.05)* indicate where transient birds exhibited significantly stronger selection for/against a given feature than residents (i.e. regardless of whether selection differed from 0 per panel a). Full model results and formula in Supplementary Table S2.

When comparing habitat selection during dispersal to residents, transient birds making large displacements expressed stronger positive selection for roads (p = 0.002) and were also more likely to select for glades (p = 0.004) than residents, but did not differ significantly from residents over any other habitat features (Fig. 3b; see Supplementary Table S2 for full model results). Similarly, during days corresponding to localized movements, transient birds again were more likely to select for roads (p = 0.011) and glades (p < 0.001) than residents, but were not different in terms of other habitat features (Fig. 3b; full results in Supplementary Table S2). Results of our supplementary models, which did not distinguish between transient modes, showed approximately the same results as our main models—i.e. residents’ selection coefficients identical, dispersers selecting for (and more strongly than residents) roads and glades and avoiding riverine areas and rugged terrain (similar to residents), with the one difference that dispersers in the aggregate were also significantly avoidant of waterbodies (RSS = 0.703; Supplementary Fig. S2, Table S3) and expressed significantly lower selection for waterbodies relative to residents (p = 0.012, Supplementary Table S4).

### Habitat Selection During Dispersal Reflects Gains in Energetic Efficiency

When travelling on roads, actively-dispersing transient birds averaged a significantly lower energetic cost of transport (CoT = 25.57 J kg^-1^ m^-1^, p < 0.001) compared to all other habitats (Fig. 4; Supplementary Table S6). Compared to the most efficient habitat type—roads—dispersers moving through acacia scrub experienced a 33.1% higher average energetic cost of transport (p < 0.001) to displace over the same net distance (50 m). They also experience significant reductions in efficiency when moving through areas of bare soil (35.9% greater CoT, p < 0.001) and glades (38.2% greater CoT, p < 0.001), with the lowest efficiency in riverine habitats (138% greater CoT, p < 0.001). However, non-dispersing residents and transient birds during local movements showed no evidence for significant differences in efficiency across habitat types (Supplementary Table S6).

**Figure 4.**
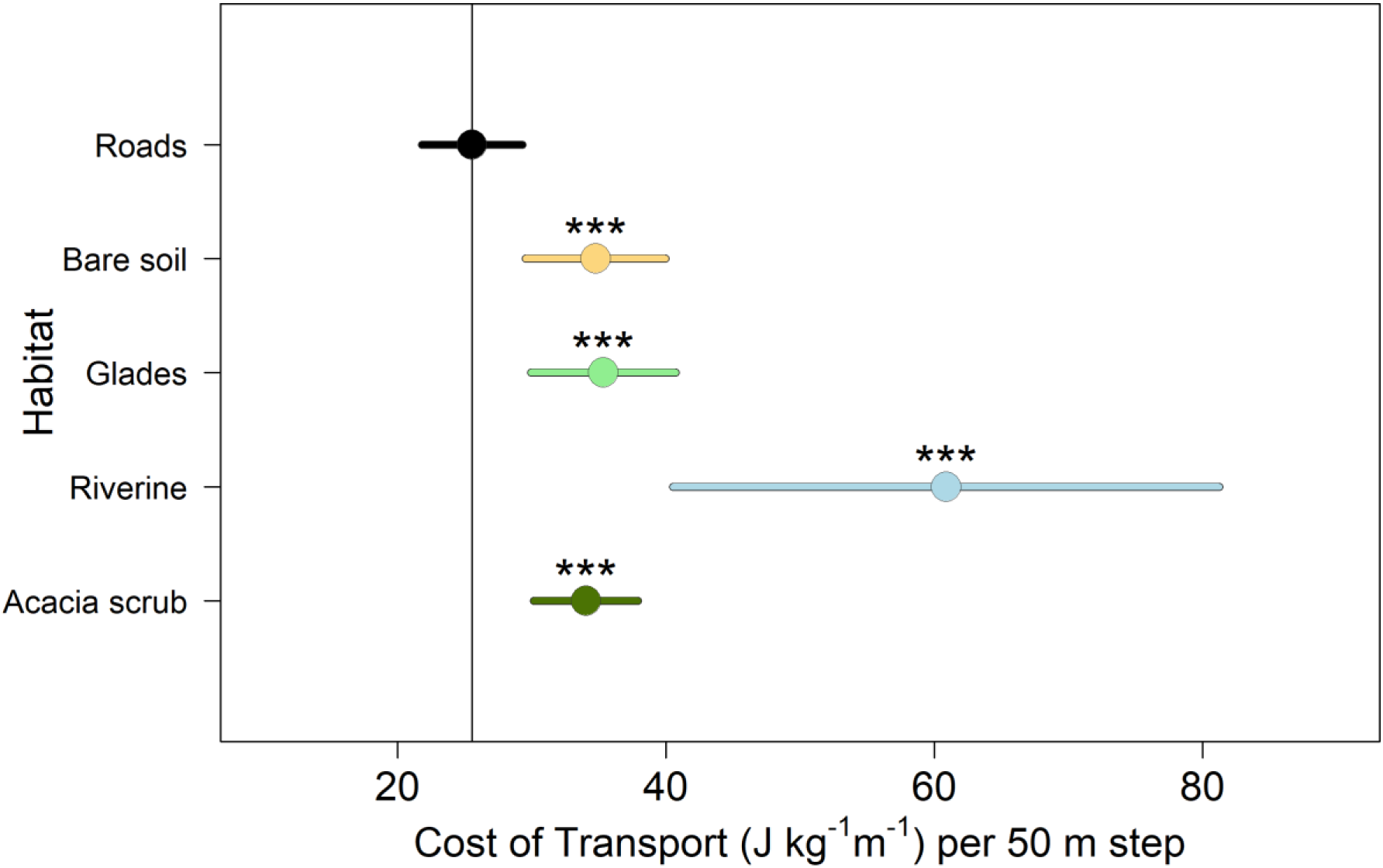
Changes in the efficiency of movement according to habitat type when making large, active dispersal movements. Circles show estimates from linear mixed-effect model (LMM) of the energetic cost of transport for actively-dispersing transient birds per 50 m net displacement across habitat type (coefficient ± 95 % confidence intervals) using roads as the reference category (intercept marked with vertical grey line), with significant increases (i.e. where movements in a given habitat were less efficient) marked by asterisks above points (*** *p<0.001*, ** *p<0.01*, \**p<0.05*). Full results, including models of efficiency for residents and local transience movements in Supplementary Table S6.

## Discussion

Our study revealed significant shifts in habitat selection by terrestrial birds during dispersal, with actively dispersing individuals selecting for open habitats (roads and glades) when making large displacements and specifically exhibiting stronger selection for these features than non-dispersing residents do. Correspondingly, we found that actively-dispersing individuals experienced a significant decrease in their cost of transport (i.e., exhibited most energetically-efficient movements) when moving along roads, whereas dispersers making local movements and residents expressed only minor differences in the cost of transport across habitat types. Our results therefore provide support for the hypothesis that changes in habitat selection expressed by terrestrial animals during dispersal could be part of a broader strategy responding to selective pressures acting on the need to achieve large displacements in an energetically efficient manner. More specifically, we found that the cost of transport was minimised (and correspondingly energetic efficiency maximised) when actively-dispersing individuals combined selection for open and linear habitats with walking in a faster, more directed manner.

In many species, the factors governing habitat selection and movement are the result of trade-offs between energetic efficiency and risk [73,74]. A number of studies in large mammals [33,75–78] have found that individuals express preferences for linear landscape features, such as roads, during periods when they make larger displacements. However, roads can also represent areas of higher disturbance and mortality risk (i.e., due to vehicle strikes), leading to a number of studies reporting avoidance of roads [79–81]. In our study, we found that transient, dispersing individuals expressed a strong positive selection for moving along roads, especially during days where they were actively displacing across the landscape, while non-dispersing residents avoided roads. This effect is especially strong when considering the relative rarity of roads in our study environment—i.e. our models suggest that actively dispersing birds should be roughly 16.6% more likely (Table S1) to make a step onto a road when roads are equally available to areas of non-road habitat, but roads actually make up less than 3% of our total study area (Table S5). At our study site (the same area as the aforementioned baboon study [33]), most roads consist of infrequently used, private dirt tracks with easily detected low-speed traffic, meaning that the mortality risks to guineafowl is quite low (in eight years of following a population of 600-1000 birds, we know of only two birds killed by vehicles).

By contrasting the features of different habitat types (Fig. S3), we could demonstrate that the potential gains in efficiency associated with roads are because the allow birds to traverse habitat in a fast and direct manner (Fig. S4). In the areas used by guineafowl, roads are generally found on slightly steeper and more variable terrain (a small effect, but different from all other habitat features except for Acacia scrub, which was slightly steeper, see Supplementary Table S7, Fig. S3).

This suggests that the importance of roads is not because they coincide with areas that have less incline or are less rugged. Birds moving along roads also did not experience different inclines relative to any other habitat (Fig. S4). Instead, we found that birds were faster and made straighter movements along roads than through all other habitat types (Supplementary Table S8., Fig. S4). Together, these results suggest that it is specifically the linearity of roads that allows individuals to traverse the landscape more directly, and thus more efficiently.

Contrary to our expectations, other open habitats (i.e. areas with bare soil or glades) were not substantially more efficient to move through than more densely vegetated areas. Nonetheless, there were interesting patterns surrounding their use by actively dispersing birds. First, the only observed steps in bare soils were made by transient birds (Supplementary Table S5) and never by non-dispersing residents (although we could not generate a robust statistical estimate of the strength of selection), likely because bare soils generally correspond to areas with little to no suitable foraging habitat. By contrast, residents showed strong positive selection for glades, which are commonly used by vulturine guineafowl for foraging [55,82]. Interestingly, the single highest selection coefficient in our models (Fig. 3, Supplementary Table S1) was for transient birds utilizing glades (they were also significantly more likely to select for glades than residents). Given the lack of energetic savings when moving on glades, the increased selection for glades by dispersing birds may instead reflect other aspects of their dispersal ecology. Dispersal in vulturine guineafowl involves individuals moving alone while trying to locate new social groups to settle into [21]. Given that the only habitats for which residents exhibited positive selection were glades, it may be the case that dispersers are selecting for glades as a means of locating groups (in addition to the higher likelihood of detecting conspecifics in the open [37,38]) in order to attempt to settle or to opportunistically forage more safely [83]. This trend is strongest during days of active dispersal movements, but was also significantly greater when making local movements relative to residents. Days of local movement correspond to periods where dispersers briefly join groups before moving on to further active displacements, and may suggest that birds are reluctant to follow groups off of glades, which is where that are most likely to forage. Together, these results suggest that it is quite likely that habitat selection by transient birds alternates between different main functions, including increasing movement efficiency on roads, increasing the chances of detecting groups on glades to potentially disperse into, and increasing foraging opportunities on glades.

In addition to changing the energetic profile, the openness of a habitat is also likely to shape predation risk. In group-living species, individuals can substantially benefit from collective detection of predators (i.e., the many eyes hypothesis [83]), which is likely to play a major role in the evolutionary ecology of vulturine guineafowl. Vulturine guineafowl are large and colourful, and live in a predator-rich environment that includes a full range of terrestrial (e.g., leopard *Panthera pardus*) and aerial (e.g., martial eagle *Polemaetus bellicosus*, Eastern chanting goshawk *Melierax poliopterus*) predators. Of these, aerial predators are likely to be particularly risky for vulturine guineafowl. In fact, the only predation event we recorded during the active phase of dispersal was by a martial eagle. The risk of aerial predators was also captured by a previous study [47] finding that dispersers avoid making large movements during the middle of the day when these predators are most active. Thus, if using open habitats introduces even more risk for individuals during the active phases of dispersal, this strengthens the support for the energy efficiency hypothesis.

Our study contributes to a growing body of evidence (e.g. [31]) on the differing habitat requirements that animals can exhibit during dispersal and substantially extends previous work by explicitly quantifying the energetic benefits that individuals can gain as function of both where and how they move when dispersing. However, making large displacements is not limited to natal dispersal. Many animals—especially those living in harsh environments—have to temporarily relocate to find resources (e.g. migration [45] or nomadism [84]), and future work may also investigate whether animals can exhibit plastic habitat preferences linked to movement under different contexts. For example, recent work has shown that migratory birds can select for different habitat features at stopover sites than in their non-migratory ranges [85], and several studies have shown how migratory ungulates shift between displacing and foraging modes in different habitats (e.g. [86,87]). As GPS tracking datasets also improve in coverage and sampling frequency, they should also open up opportunities to investigate movement strategies that could be expressed by resident individuals. For example, many animals in hot regions are more active (foraging, moving) in the mornings and evenings. The limited time available for resource acquisition could drive strategies to increase the speed of movement—such as the ‘morning commute’ of baboons following roads to reach distant foraging areas before the day heats up [33]. Thus, our study approach could be replicated by looking at changes in habitat selection and energetic efficiency across different hours of the day.

Finally, we note that in our study we were limited to making inferences about the selection for more energetically efficient movements in two separate analytical steps. This is because dispersers moved through areas in which we had little to no prior GPS data (unlike areas used by their natal groups), thereby limiting our ability to generate an ‘efficiency landscape’. While we could have made some inferences about the cost of transport for different habitats, our analysis would have been based on fitting measured cost of transport values in animals’ chosen steps and inferred values for alternative steps. However, future work that is focused more closely on areas with high GPS coverage, for example by looking at changing selection patterns over the course of the day, could use repeated observations of movements by individuals over the same areas to produce a cost of transport layer, and directly test whether individuals are selecting for habitats with lower cost of transport when making larger displacements. We anticipate that such studies will provide rich grounds for gaining insights into the process of individual (or collective) decision-making by animals on the move.

## Acknowledgements

We thank the Mpala Research Centre, the Kenyan Wildlife Service, and the Ornithological Section of the National Museums of Kenya for supporting this research work. We are especially grateful to Dr. Peter Njoroge from the NMK for the long-term collaboration on our project, and Dr. Fred Omengo for his continued advice and oversight on behalf of the KWS. We also thank the Ol Jogi Wildlife Conservancy, Mr. Peter Jessel, and El Karama Ranch for their support in allowing us to track dispersing guineafowl on their properties. We are grateful to Wismer Cherono, John Wanjala, Monicah Wambui, John Ewoi, Martha Sisto, and Danai Papageorgiou for their contributions to data collection and field assistance.

## Ethical statement

Work was conducted under a Research Authorisation (KWS-0016-01-21) and a Capture Permit issued by the Kenyan Wildlife Service, research authorisations from the Wildlife Research and Training Institute, and in affiliation with the National Museums of Kenya. Further research permits were granted by the Max Planck Society Ethikrat Committee (2016_13/1), the National Environment Management Authority (NEMA/AGR/68/2017) and the National Commission for Science, Technology and Innovation of Kenya (NACOSTI/P/16/3706/6465, NACOSTI/P/21/8699).

