## Supplementary Materials for "Habitat selection during dispersal reduces the energetic cost of transport when making large displacements"

### SUPPLEMENTARY MATERIAL

Dispersal Data

We recorded a total of 10,357,429 GPS fixes (across two temporal sampling resolutions) from 32 individuals, from which we extracted 61,435 5-minute displacements of at least 10m (i.e., our observed steps). The majority of the steps in our dataset came from non-dispersing residents (57.9%), with the remaining steps from transient birds divided into periods of large, active dispersal movements (12.0%) and more-local transient movements (30.1%). Generating 20 alternative steps for each observed step resulted in a total of 1,198,789 5-minute steps, of which we removed 5,793 (279 observed steps and 5,514 alternate) where no value could be assigned to one of the environmental covariates at the endpoint of the step (empty data), leaving 1,192,996 steps (61,156 observed) used for the Step-Selection Model. The median step length for a 5-minute (observed) step was 28.41 meters.

**Supplementary Table S1.** Results of a step selection model investigating the absolute strength of selection for different habitat features within different movement stages (i.e. whether individuals in a given movement stage select for or against a given habitat). For binomial raster layers (Waterbodies or Roads), reference level was the absence of the given habitat feature. For categorical rasters (Habitat Cover type), Acacia scrub was set as the reference level as it was the most prevalent and abutted all other cover types. Terrain ruggedness was a continuous variable. Selection coefficient values represent the estimated difference in log(RSS) values between given feature and the reference level among individuals at a given stage. Significant values (given in bold) show where the strength of selection was significantly positive or negative within a given stage. NA values indicate no steps assigned to a given stage were assigned to a given habitat covariate.

**Habitat selection coefficients by stage—Resident, Transient (local), or Transient (large displacements).**

$n_{total}$ = 1,181,263 steps (observed + alternate), $n_{observed}$ = 60,657steps

**Formula:** *Step* [observed vs alternate] ~ *Waterbodies:stage +Roads:stage + Ruggedness:stage + Cover type:stage + stepID*

| Stage: Covariate | Sel. Coef. | exp(Coef) | se(Coef) | z | Pr(>\|z\|) |
|---|---|---|---|---|---|
| Res:Waterbody | 0.086 | 1.090 | 0.069 | 1.241 | 0.214 |
| TransLoc:Waterbody | -0.207 | 0.813 | 0.164 | -1.264 | 0.206 |
| TransDis:Waterbody | -0.292 | 0.747 | 0.186 | -1.571 | 0.116 |
| **Res:Roads** | **-0.057** | **0.944** | **0.026** | **-2.212** | **0.027** |
| TransLoc:Roads | 0.062 | 0.064 | 0.039 | 1.562 | 0.118 |
| **TransDis:Roads** | **0.150** | **1.162** | **0.062** | **2.406** | **0.016** |
| **Res:Glades** | **0.327** | **1.387** | **0.021** | **15.68** | **<0.001** |
| **TransLoc:Glades** | **0.495** | **1.941** | **0.027** | **18.58** | **<0.001** |
| **TransDis:Glades** | **0.521** | **1.684** | **0.051** | **10.16** | **<0.001** |
| **Res:Riverine** | **-0.252** | **0.778** | **0.059** | **-4.230** | **<0.001** |
| **TransLoc:Riverine** | **-0.290** | **0.748** | **0.125** | **-2.315** | **0.021** |
| **TransDis:Riverine** | **-0.441** | **0.643** | **0.134** | **-3.289** | **0.001** |
| **Res:Ruggedness** | **-0.104** | **0.901** | **0.014** | **-7.252** | **<0.001** |
| **TransLoc:Ruggedness** | **-0.076** | **0.927** | **0.019** | **-4.087** | **<0.001** |
| **TransDis:Ruggedness** | **-0.096** | **0.908** | **0.024** | **-3.992** | **<0.001** |

Concordance = 0.523, se = 0.001

Likelihood ratio test = 861 on 15 df, p < 2 e-16

Wald test = 855 on 15 df, p < 2 e-16

Score (logrank) test = 859 on 15 df, p < 2 e-16

Stages: *TransDis* = actively-dispersing transient birds on days with large displacements, *TransLoc* = transient birds during days of local movement, *Res* = non-dispersing residents.

**Supplementary Table S2.** Results of a step selection model investigating how the strength of selection for various habitat features changes as a function of movement stages (i.e. whether the selection behaviours of actively-dispersing birds are significantly different than those of non-dispersing residents). Non-dispersing residents' movements were used as the reference category against which two dispersal modes (large displacements and local movements) were compared. For binomial raster layers (Waterbodies or Roads), reference level was the absence of the given habitat feature. For categorical rasters (Habitat Cover type), Acacia scrub was set as the reference level as it was the most prevalent and abutted all other cover types. Terrain ruggedness was a continuous variable. Selection coefficient values represent the estimated difference in log(RSS) values for a given feature relative to residents' selection coefficient (also given here). Significant values (given in bold) show where dispersing birds' strength of selection was significantly different from that of non-dispersing residents. Note that the same stage:feature coefficient estimates found in Table S2 can be produced by summing a given coefficient here with its reference value (e.g. Res:Glades + TransDis:Glades in this table is equal to TransDis:Glades in S2), but that the estimates of significance are based on different comparisons. NA values indicate no comparative estimate could be produced relative to residents.

**Habitat selection coefficients across stages—Resident, Transient (local), or Transient (large displacements).**

$n_{total}$ = 1,181,263 steps (observed + alternate), $n_{observed}$ = 60,657steps

**Formula:** *Step* [observed vs alternate] ~ Waterbodies + Roads + Ruggedness + Cover type + *Waterbodies:stage +Roads:stage + Ruggedness:stage + Cover type:stage + stepID*

| Stage: Covariate | Sel. Coef. | exp(Coef) | se(Coef) | z | Pr(>\|z\|) |
|---|---|---|---|---|---|
| Res:Waterbody | 0.086 | 1.090 | 0.069 | 1.241 | 0.214 |
| TransLoc:Waterbody | -0.293 | 0.746 | 0.178 | -1.647 | 0.099 |
| TransDis:Waterbody | -0.378 | 0.685 | 0.198 | -1.905 | 0.057 |
| **Res:Roads** | **-0.057** | **0.944** | **0.026** | **-2.212** | **0.027** |
| **TransLoc:Roads** | **0.119** | **1.126** | **0.047** | **2.520** | **0.012** |
| **TransDis:Roads** | **0.207** | **1.230** | **0.067** | **3.072** | **0.002** |
| **Res:Glades** | **0.327** | **1.387** | **0.021** | **15.68** | **<0.001** |
| **TransLoc:Glades** | **0.168** | **1.183** | **0.034** | **4.970** | **<0.001** |
| **TransDis:Glades** | **0.194** | **1.214** | **0.055** | **3.505** | **<0.001** |
| **Res:Riverine** | **-0.252** | **0.778** | **0.059** | **-4.230** | **<0.001** |
| TransLoc:Riverine | -0.039 | 0.962 | 0.139 | -0.280 | 0.780 |
| TransDis:Riverine | -0.190 | 0.827 | 0.147 | -1.293 | 0.196 |
| **Res:Ruggedness** | **-0.104** | **0.901** | **0.014** | **-7.252** | **<0.001** |
| TransLoc:Ruggedness | 0.028 | 1.028 | 0.023 | 1.193 | 0.233 |
| TransDis:Ruggedness | 0.008 | 1.008 | 0.028 | 0.268 | 0.789 |

Concordance = 0.523, se = 0.001

Likelihood ratio test = 861 on 15 df, p < 2 e-16

Wald test = 855 on 15 df, p < 2 e-16

Score (logrank) test = 859 on 15 df, p < 2 e-16

Stages: *TransDis* = actively-dispersing transient birds on days with large displacements, *TransLoc* = transient birds during days of local movement, *Res* = non-dispersing residents.

**Supplementary Table S3.** Results of a step selection model investigating habitat preferences as a function of whether and individual was dispersing or not (i.e. irrespective of daily movement mode for transient dispersers). For binomial raster layers (Waterbodies or Roads), reference level was the absence of the given habitat feature. For categorical rasters (Habitat Cover type), Acacia scrub was set as the reference level as it was the most prevalent and abutted all other cover types. Terrain ruggedness was a continuous variable. Significant values given in bold. NA values indicate no steps assigned to a given stage were assigned to a given habitat covariate.

**Habitat selection coefficients by stage—Resident or Transient**

$n_{total}$ = 1,179,950 steps (observed + alternate), $n_{observed}$ = 60,657steps

**Formula:** *Step* [observed vs alternate] ~ *Waterbodies:stage +Roads:stage + Cover type:stage + Ruggedness:stage + stepID*

| Stage: Covariate | Sel. Coef. | exp(Coef) | se(Coef) | z | Pr(>\|z\|) |
|---|---|---|---|---|---|
| Resident:Waterbody | 0.086 | 1.090 | 0.069 | 1.241 | 0.214 |
| **Transient:Waterbody** | **-0.247** | **0.781** | **0.123** | **-2.011** | **0.044** |
| **Resident:Roads** | **-0.057** | **0.944** | **0.026** | **-2.212** | **0.027** |
| **Transient:Roads** | **0.087** | **10.91** | **0.033** | **2.560** | **0.009** |
| **Resident:Glades** | **0.327** | **1.387** | **0.021** | **15.68** | **<0.001** |
| **Transient:Glades** | **0.501** | **1.650** | **0.024** | **21.17** | **<0.001** |
| **Resident:Riverine** | **-0.252** | **0.778** | **0.059** | **-4.230** | **<0.001** |
| **Transient:Riverine** | **-0.365** | **0.694** | **0.091** | **-3.993** | **<0.001** |
| **Resident:Ruggedness** | **-0.104** | **0.901** | **0.014** | **-7.252** | **<0.001** |
| **Transient:Ruggedness** | **-0.084** | **0.920** | **0.015** | **-5.700** | **<0.001** |

Concordance = 0.523, se = 0.001

Likelihood ratio test = 857 on 10 df, p < 2 e-16

Wald test = 852 on 10 df, p < 2 e-16

Score (logrank) test = 855 on 10 df, p < 2 e-16

**Supplementary Table S4.** Results of a step selection model investigating how the strength of selection for various habitat features changes as a function of movement stages (i.e. whether the selection behaviours of actively-dispersing birds are significantly different than those of non-dispersing residents). Non-dispersing residents' movements were used as the reference category against which dispersal movements (irrespective of daily movement mode) were compared. For binomial raster layers (Waterbodies or Roads), reference level was the absence of the given habitat feature. For categorical rasters (Habitat Cover type), Acacia scrub was set as the reference level as it was the most prevalent and abutted all other cover types. Terrain ruggedness was a continuous variable. Selection coefficient values represent the estimated difference in log(RSS) values for a given feature relative to residents' selection coefficient (also given here). Significant values (given in bold) show where dispersing birds' strength of selection was significantly different from that of non-dispersing residents. Note that the same stage:feature coefficient estimates found in Table S4 can be produced by summing a given coefficient here with its reference value (e.g. Resident:Glades + Transient:Glades in this table is equal to Transient:Glades in S4), but that the estimates of significance are based on different comparisons. NA values indicate no comparative estimate could be produced relative to residents.

**Habitat selection coefficients across stages—Resident or Transient**

$n_{total}$ = 1,179,950 steps (observed + alternate), $n_{observed}$ = 60,657steps

**Formula:** *Step* [observed vs alternate] ] ~ Waterbodies + Roads + Ruggedness + Cover type + *Waterbodies:stage +Roads:stage + Cover type:stage + Ruggedness:stage + stepID*

| Stage: Covariate | Sel. Coef. | exp(Coef) | se(Coef) | z | Pr(>\|z\|) |
|---|---|---|---|---|---|
| Resident:Waterbody | 0.086 | 1.090 | 0.069 | 1.241 | 0.214 |
| **Transient:Waterbody** | **-0.353332** | **0.717** | **0.141** | **-2.362** | **0.018** |
| **Resident:Roads** | **-0.057** | **0.944** | **0.026** | **-2.212** | **0.027** |
| **Transient:Roads** | **0.144** | **1.155** | **0.042** | **3.410** | **<0.001** |
| **Resident:Glades** | **0.327** | **1.387** | **0.021** | **15.68** | **<0.001** |
| **Transient:Glades** | **0.174** | **1.190** | **0.032** | **5.512** | **<0.001** |
| **Resident:Riverine** | **-0.252** | **0.778** | **0.059** | **-4.230** | **<0.001** |
| Transient:Riverine | -0.113 | 0.893 | 0.109 | -1.039 | 0.299 |
| **Resident:Ruggedness** | **-0.104** | **0.901** | **0.014** | **-7.252** | **<0.001** |
| Transient:Ruggedness | 0.020 | 1.020 | 0.021 | 0.976 | 0.329 |

Concordance = 0.523, se = 0.001

Likelihood ratio test = 857 on 10 df, p < 2 e-16

Wald test = 852 on 10 df, p < 2 e-16

Score (logrank) test = 855 on 10 df, p < 2 e-16

**Supplementary Table S5.** Proportion of habitat available (i.e. proportion of cells in a given raster layer attributed to a given habitat feature) and the proportion of observed steps attributed to each habitat cover type or feature for each movement category (i.e. actively-dispersing transient birds on days with large displacements, transient birds during days of local movement, and non-dispersing residents).

| | **Proportional use of each habitat cover type** | | | | |
|---|---|---|---|---|---|
| | **Acacia Scrub** | **Glades** | **Riverine** | **Bare Soil** | **Black Cotton** |
| **Proportion available** | 0.724 | 0.012 | 0.021 | 0.051 | 0.191 |
| **Transient (large movements)** | 0.774 | 0.158 | 0.015 | 0.026 | 0.027 |
| **Transient (local)** | 0.654 | 0.332 | 0.008 | 0.005 | 0.001 |
| **Residents** | 0.727 | 0.247 | 0.026 | 0 | 0 |
| | **Proportional use of habitat features** | | | | |
| | **Roads** | **Waterbodies** | | | |
| **Proportion available** | 0.027 | 0.011 | | | |
| **Transient (large movements)** | 0.053 | 0.007 | | | |
| **Transient (local)** | 0.055 | 0.004 | | | |
| **Residents** | 0.072 | 0.013 | | | |

**Supplementary Table S6.** Results of 3 linear models fit to the total energetic cost of transport, in Joules $kg^{-1} m^{-1}$, over a 50-meter net displacement and its response to habitat type, with each model drawing from movements expressed by individuals within a given movement stage. Reference level of habitat is roads in all models.

| **Energetic cost of transport per 50m net displacement by movement stage** | | | | |
|---|---|---|---|---|
| **Model 1: Non-dispersing residents**<br>n = 359 displacements<br>*Cost of transport$_{(Resident)}$ ~ Habitat* | | | | |
| | **Estimate** | **Std. Error** | **t value** | **Pr(>\|t\|)** |
| **Intercept (Roads)** | 30.376 | 4.964 | 6.119 | **<0.001** |
| **Habitat: Glades** | 4.024 | 5.480 | 0.734 | 0.463 |
| **Habitat: Riverine** | 7.482 | 11.817 | 0.633 | 0.527 |
| **Habitat: Acacia scrub** | 6.468 | 5.088 | 1.271 | 0.204 |
| **Model 2: Transient birds, local movements**<br>n = 512 displacements<br>*Cost of transport$_{(Transient-Local)}$ ~ Habitat* | | | | |
| | **Estimate** | **Std. Error** | **t value** | **Pr(>\|t\|)** |
| **Intercept (Roads)** | 31.397 | 3.648 | 8.606 | **<0.001** |
| **Habitat: Glades** | 3.197 | 4.333 | 0.738 | 0.461 |
| **Habitat: Riverine** | 9.048 | 8.611 | 1.051 | 0.294 |
| **Habitat: Acacia scrub** | 5.735 | 3.793 | 1.512 | 0.131 |
| **Model 3: Transient birds, large movements**<br>n = 930 displacements<br>*Cost of transport$_{(Transient-Large)}$ ~ Habitat* | | | | |
| | **Estimate** | **Std. Error** | **t value** | **Pr(>\|t\|)** |
| **Intercept (Roads)** | 26.253 | 2.080 | 12.623 | **<0.001** |
| **Habitat: Bare Soil** | 9.169 | 2.893 | 3.170 | **0.002** |
| **Habitat: Glades** | 11.023 | 2.996 | 3.679 | **<0.001** |
| **Habitat: Riverine** | 38.601 | 11.296 | 3.417 | **<0.001** |
| **Habitat: Acacia scrub** | 9.020 | 2.157 | 4.182 | **<0.001** |

**Supplementary Table S7.** Results of 2 linear models investigating the relationship between habitat features (roads and cover types) and underlying terrain topography—either (1) terrain slope, or (2) terrain ruggedness (using the terrain ruggedness index, TRI). Each datapoint (n = 489142) corresponds to each unique cell in the habitat cover raster which contained an observed or alternative step (i.e. all cells which contributed to our step-selection analysis). Where cells were assigned to roads in addition to another feature (due to separate raster layer for roads), only the road label was used. Reference level of habitat is roads in all models.

<table>
<tr><th colspan="5">Model 1: Terrain slope across habitat features<br>n = 489142 cells<br>Slope (degrees) ~ Habitat</th></tr>
<tr><th></th><th>Estimate</th><th>Std. Error</th><th>t value</th><th>Pr(>|t|)</th></tr>
<tr><td>Intercept (Roads)</td><td>3.364</td><td>0.011</td><td>298.8</td><td><0.001</td></tr>
<tr><td>Habitat: Black cotton soil</td><td>0.504</td><td>0.012</td><td>43.27</td><td><0.001</td></tr>
<tr><td>Habitat: Glades</td><td>-1.402</td><td>0.031</td><td>-45.05</td><td><0.001</td></tr>
<tr><td>Habitat: Bare soil</td><td>-0.222</td><td>0.013</td><td>-16.46</td><td><0.001</td></tr>
<tr><td>Habitat: Acacia scrub</td><td>-0.881</td><td>0.028</td><td>-31.99</td><td><0.001</td></tr>
<tr><td>Habitat: Riverine</td><td>-0.823</td><td>0.021</td><td>-39.08</td><td><0.001</td></tr>
</table>

<table>
<tr><th colspan="5">Model 2: Terrain ruggedness across habitat features<br>n = 489142 cells<br>Ruggedness (TRI) ~ Habitat</th></tr>
<tr><th></th><th>Estimate</th><th>Std. Error</th><th>t value</th><th>Pr(>|t|)</th></tr>
<tr><td>Intercept (Roads)</td><td>1.505</td><td>0.005</td><td>333.1</td><td><0.001</td></tr>
<tr><td>Habitat: Black cotton soil</td><td>0.179</td><td>0.005</td><td>38.38</td><td><0.001</td></tr>
<tr><td>Habitat: Glades</td><td>-0.522</td><td>0.012</td><td>-41.80</td><td><0.001</td></tr>
<tr><td>Habitat: Bare soil</td><td>-0.069</td><td>0.005</td><td>-12.81</td><td><0.001</td></tr>
<tr><td>Habitat: Acacia scrub</td><td>-0.342</td><td>0.011</td><td>-30.94</td><td><0.001</td></tr>
<tr><td>Habitat: Riverine</td><td>-0.249</td><td>0.008</td><td>-29.43</td><td><0.001</td></tr>
</table>

**Supplementary Table S8.** Results of 3 models investigating the relationship between habitat features (roads and cover types) and behavioural aspects of movement—speed, path straightness, and the incline that birds moved over—during each of the 50-meter net displacements used in our energetic analyses (n=1801 observations). Movement speed (Model 1) was modelled as a linear model fit to the log-transformed mean speed-while-moving (i.e. the mean of all speed values in m $s^{-1}$ from all points assigned to HMM states 2-4 within a given 50m displacement) as the response variable. Path straightness (Model 2) was modelled by fitting a beta regression to straightness index values (SI, net displacement/cumulative path length therein) for each net displacement. Movement incline (3) was modelled with a linear model fit to the incline (percentile, absolute net elevation change/net displacement distance) that birds moved over in the course of each displacement. Reference level of habitat is roads in all models.

**Movement characteristics during 50m net displacements**

**Model 1: Movement speed**
*log(mean(speed-while-moving)) ~ Habitat*

| | **Estimate** | **Std. Error** | **t value** | **Pr(>\|t\|)** |
|---|---|---|---|---|
| **Intercept (Roads)** | -0.315 | 0.070 | -4.527 | **<0.001** |
| **Habitat: Bare soil** | -0.310 | 0.114 | -2.712 | **0.007** |
| **Habitat: Glades** | -0.313 | 0.086 | -3.640 | **<0.001** |
| **Habitat: Riverine** | -0.703 | 0.215 | -3.264 | **0.001** |
| **Habitat: Acacia scrub** | -0.359 | 0.072 | -4.978 | **<0.001** |

**Model 2: Path straightness**
*SI ~ Habitat, family=beta*

| | **Estimate** | **Std. Error** | **t value** | **Pr(>\|t\|)** |
|---|---|---|---|---|
| **Intercept (Roads)** | 1.587 | 0.081 | 19.521 | **<0.001** |
| **Habitat: Bare soil** | 0.213 | 0.130 | -1.639 | 0.101 |
| **Habitat: Glades** | -0.282 | 0.098 | -2.867 | **0.004** |
| **Habitat: Riverine** | -0.563 | 0.239 | -2.355 | **0.019** |
| **Habitat: Acacia scrub** | -0.279 | 0.083 | -3.350 | **<0.001** |

**Model 3: Displacement incline**
*Incline(%) ~ Habitat*

| | **Estimate** | **Std. Error** | **t value** | **Pr(>\|t\|)** |
|---|---|---|---|---|
| **Intercept (Roads)** | 3.950 | 0.327 | 12.096 | **<0.001** |
| **Habitat: Bare soil** | -0.152 | 0.535 | -0.284 | 0.777 |
| **Habitat: Glades** | -0.518 | 0.404 | -1.284 | 0.199 |
| **Habitat: Riverine** | -0.321 | 1.011 | -0.317 | 0.751 |

| | | | | |
|---|---|---|---|---|
| **Habitat: Acacia scrub** | 0.257 | 0.338 | 0.760 | 0.447 |

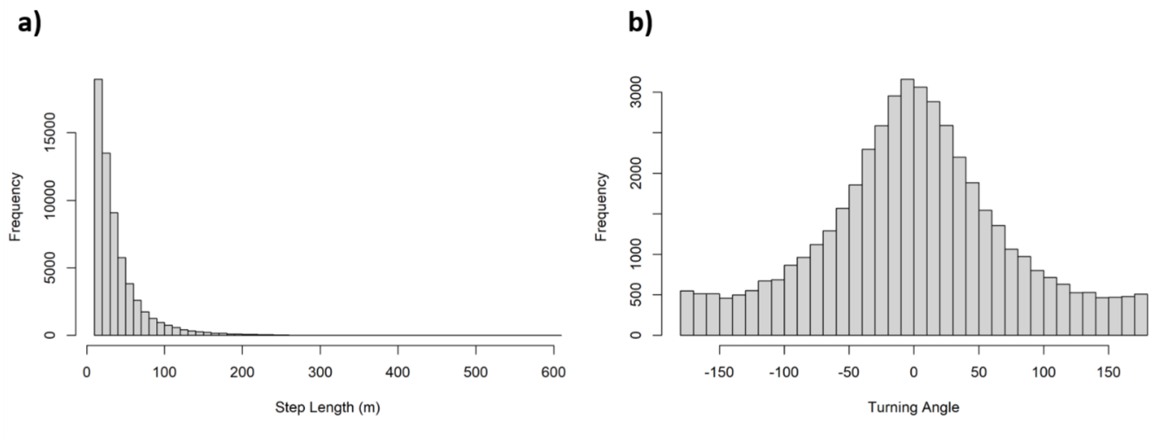


**Supplementary Figure S1. Population-level distributions of step lengths (A) and turning angles (B) from 61,453 observed steps. Step selection models were fit using the distributions for each of 32 individuals.**

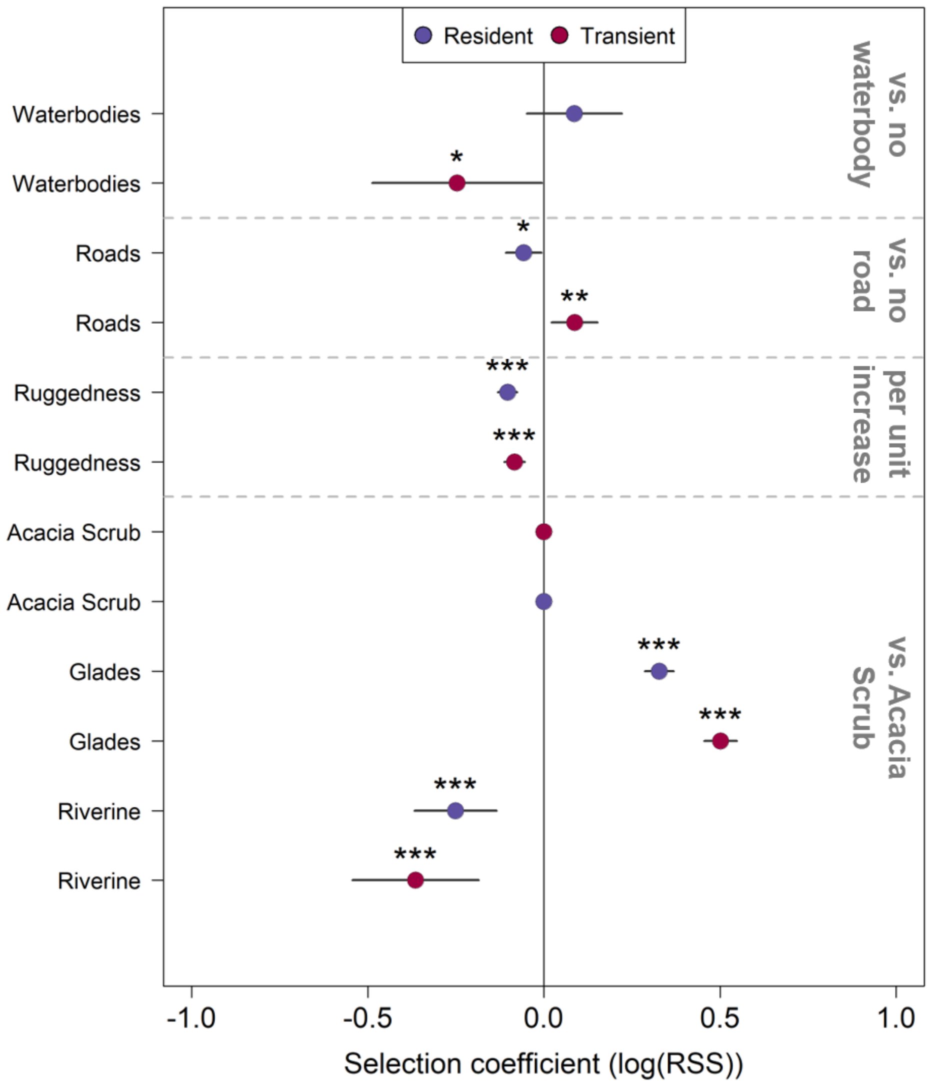


**Supplementary Figure S2: Estimated selection coefficients for each habitat covariate for dispersing (transient) and non-dispersing (resident) individuals.** Points represent log-transformed estimated relative selection strength (RSS) values from our step-selection analysis. Black bars 95% confidence intervals, indicating strength of habitat selection preferences for (where log(RSS) > 0) or avoidance of (where log(RSS) < 0) a given habitat type. Data correspond to movements of transient dispersers (irrespective of daily movement mode, red) and non-dispersing residents (purple). Dashed grey line separate variables from different raster layers. Grey text (right) indicates reference level against which selection coefficients are contrasted. Stars above a point indicate the statistical significance (*** *p<0.001*, ** *p<0.01*, **p<0.05)* of a given estimate.

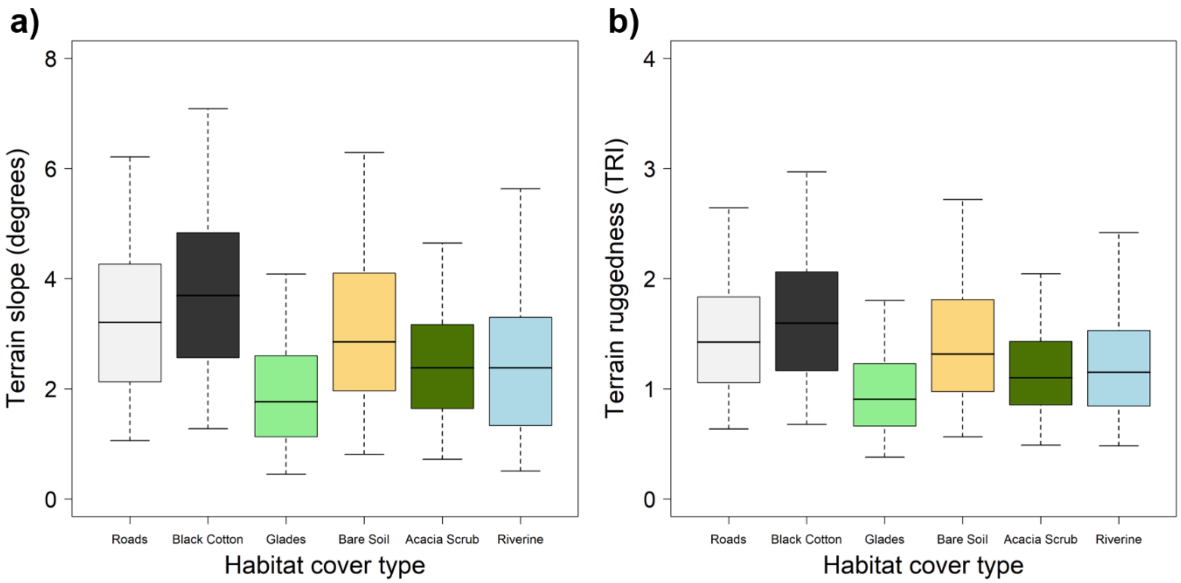


**Supplementary Figure S3. Terrain slope and ruggedness associated with different habitat features across all sampled cells within our study area.** Panels show box-and-whisker plots for terrain slope (a) and ruggedness (b) among all unique raster cells which contained either an observed or generated step in our 5-minute data used to parameterize the step-selection analysis. Boxes show the median value, bounded by the lower and upper quartiles, while whiskers show the 95$^{th}$ percentile ranges (0.05 and 0.95 quantiles respectively) for each value according to habitat cover type. Cells corresponding to both roads and another cover type (due to labels coming from separate raster layers) were considered only as roads here. Full model results in Supplementary Table S7.

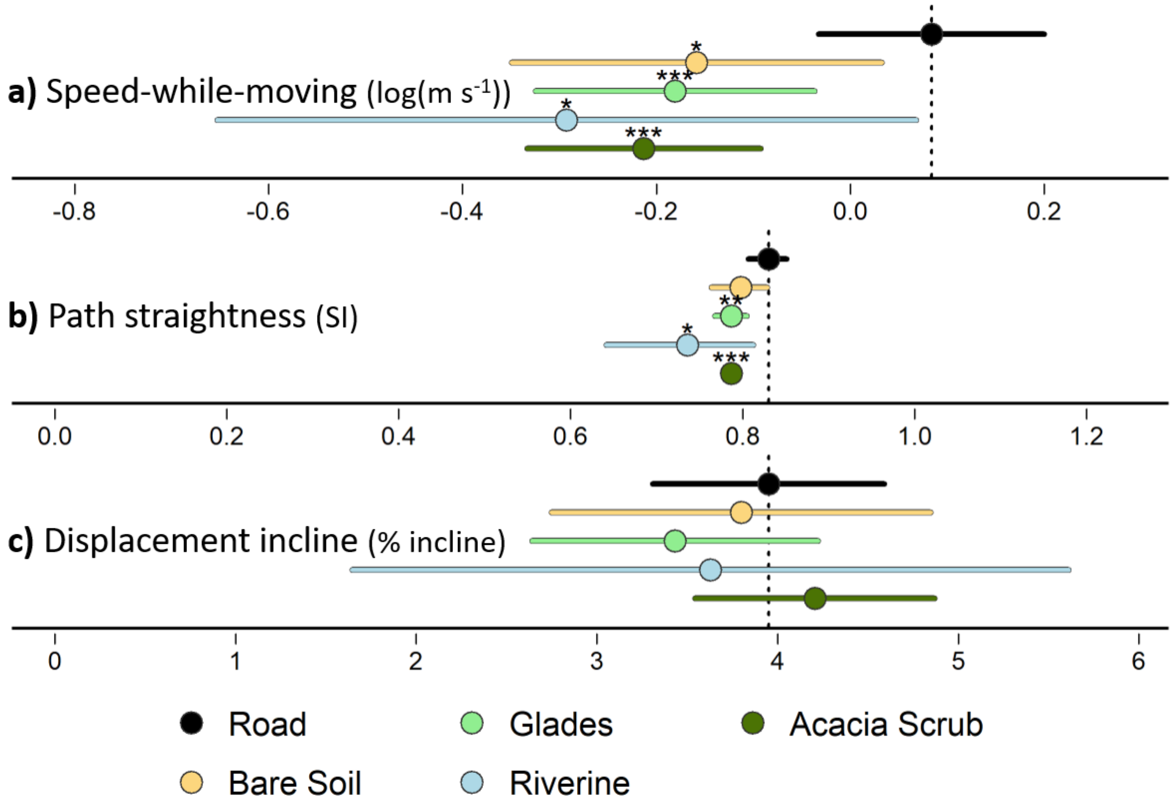


**Supplementary Figure S4. Behavioural features of movement according to habitat type over 50-meter net displacements.** (a–c) Summaries of regression models characterizing (coefficient ± 95% confidence intervals) movement behaviours expressed during 50-meter net displacements within a given habitat. All models used movements of individuals along roads as the reference category (dotted vertical lines correspond to model intercept), with significant differences (*** $p<0.001$, ** $p<0.01$, * $p<0.05$) marked by asterisks above points. Speed-while-moving (a) was calculated as the average speed at all moving (HMM states 2-4) points within a given net displacement. Path straightness (b) was calculated as a straightness index (SI) based on the ratio of the net displacement distance (approx. 50m) to the sum of all steps taken to reach that net displacement. Displacement incline (c) was calculated based on the ratio of horizontal net displacement (approx. 50m) to the absolute net change in elevation between the first and last points of a displacement. Full model results in Supplementary Table S8.
